# Stochastic dynamics of extra-chromosomal DNA

**DOI:** 10.1101/2019.12.15.876714

**Authors:** Yuriy Pichugin, Weini Huang, Benjamin Werner

**Affiliations:** Department of Evolutionary Theory, Max Planck Institute for Evolutionary Biology, Plön, Germany; Department of Mathematics, Queen Mary University of London, United Kingdom; Group of Theoretical Biology, The State Key Laboratory of Biocontrol, School of Life Science, Sun Yat-sen University, Guangzhou, 510060 China; Evolutionary Dynamics Group, Centre for Cancer Genomics and Computational Biology, Barts Cancer Centre, Queen Mary University of London, Charterhouse Square, London, United Kingdom EC1M 6BQ

## Abstract

Nuclear extra-chromosomal DNA (ecDNA) is highly prevalent in human tumours. ecDNA amplifications can promote accessible chromatin, oncogen over-expression and imply a worse clinical prognosis. Yet, little is known about the evolutionary process of ecDNA in human cancers. Here, we develop the theoretical foundation of the ecDNA somatic evolutionary process combining mathematical, computational and experimental approaches. We show that random ecDNA segregation leads to unintuitive dynamics even if ecDNA is under very strong positive selection. Patterns of inter- and intra-tumour ecDNA and chromosomal heterogeneity differ markedly and standard approaches are not directly applicable to quantify ecDNA evolution. We show that evolutionary informed modelling leads to testable predictions on how to distinguish positively selected from neutral ecDNA dynamics. Our predictions describe the dynamics of circular amplicons in GBM39 cell line experiments, suggesting a 300% fitness increase due to circular extra-chromosomal *EGFRvIII* amplifications. Our work lies the basis for further studies to quantitate ecDNA somatic evolutionary processes.

## Introduction

Recent genome profiling studies of both healthy and cancerous human tissues found a high prevalence of nuclear extra-chromosomal DNA (ecDNA) [1, 2, 3, 4]. ecDNA occurs during episodes of chromosomal instability and contributes to cancer initiation and treatment resistance [5, 6, 7, 8, 9]. Double minute oncogene amplifications exist in 31.7% of all neuroblastoma and highly amplified extra-chromosomal KRAS copies appear at relapse of anti-EGFR targeted therapies [1, 4, 9]. Yet, the evolutionary fade of specific ecDNA elements within individual tumours remains unknown. It is of great interest and practical need to develop methods to quantify ecDNA driven somatic evolutionary processes.

One major complication is the unique nature of the ecDNA somatic evolutionary process. Circular ecDNA lacks centromeres, causing random segregation of ecDNA into daughter cells after division [5] (Figure 1). Compared to somatic changes of chromosomal DNA, this adds additional stochasticity to the evolutionary dynamics of ecDNA [10, 11, 12]. Consequently, patterns of inter- and intra-patient ecDNA heterogeneity differ compared to chromosomal point mutations [13, 14, 15]. Methods that quantify somatic evolutionary processes based on chromosomal point mutations, such as phylogenetic trees [16, 17, 18, 19, 20, 21], ratios of synonymous and non-synonymous mutations (dN/dS) [22, 23, 24], or site-frequency spectra [25, 26, 27, 10, 28, 29] are not directly applicable to quantify the evolutionary dynamics of ecDNA.

**Figure 1.**
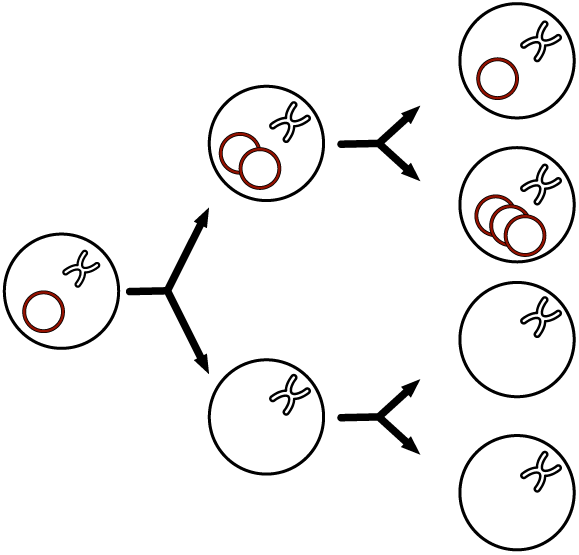
Schematics of extra-chromosomal DNA (ecDNA) population dynamics. ecDNA (circels) is randomly segregated between daughter cells. The resulting dynamics of ecDNA is markedly different compared to chromosomal alterations. For example, a cell can lose all ecDNA by chance, whereas other cells can acquire many copies of ecDNA. A growing cell population naturally becomes a mix of cells with and without ecDNA. Here, we describe how the inter- and intra-tumour ecDNA heterogeneity resembles the underlying evolutionary dynamic process, e.g. how one may quantify fitness consequences of ecDNA.

Here, we discuss the stochastic dynamics of ecDNA in a growing tumour population. We show that the random segregation of ecDNA has unintuitive consequences. For example, the frequency of cells with ecDNA can intermittently drop, even if ecDNA conveys a high selective advantage. We present a mathematical and computational analysis of the expected inter- and intra-tumour ecDNA heterogeneity under different fitness scenarios and discuss strategies to infer these scenarios from cancer genomic data. We show that a single time point measurement at single cell resolution can be sufficient to disentangle different somatic evolutionary scenarios. Furthermore, we compare the theoretically predicted ecDNA heterogeneity patterns with experimental observations. Growth experiments with the glioblastoma cell line GBM39 are consistent with very strong positive selection and an approximate fitness increase of 300% due to circular extra-chromosomal *EGFRvIII* amplifications. In summary, our work describes the stochastic population dynamics of ecDNA in a growing tumour and provides a framework to quantify ecDNA fitness effects.

## Results

In the following, we integrate a mathematical analysis of the ecDNA somatic evolutionary process, individual based stochastic computer simulations and *in vitro* ecDNA heterogeneity patterns from growth experiments of the glioblastoma cell line GBM39 with the known circular extra-chromosomal *EGFRvIII* amplification [1, 9]. For a better grasp on some of the peculiar properties of ecDNA evolution, we first discuss deterministic ecDNA dynamics. One important distinct future of the ecDNA evolutionary process is the random inheritance of ecDNA copies into daughter cells (Figure 1). The probability for a daughter cell to carry *k* ecDNA copies upon proliferation is given by a Binomial distribution with success probability 1/2 (Figure 1 and Methods). This implies that one daughter cell might not inherit any ecDNA by chance, which is distinctively different to the inheritance of somatic chromosomal point mutations. Assuming a cell carries *k* copies of ecDNA, a daughter cell would not inherit ecDNA with probability 2^−2*k*^. In other words, if a cell carries many copies of ecDNA, a daughter cell without ecDNA is unlikely. However, if a cell carries a single ecDNA copy, the subsequent probability for a daughter cell without ecDNA is 1/4. We therefore should expect a tumour to be a mix of cells with and without ecDNA. Our results show that the exact tumour composition of cells with and without ecDNA is stochastic especially at the early stages of tumour evolution and critically depends on the underlying ecDNA fitness distribution.

### Frequencies of cells with and without ecDNA

We now turn to the expected frequencies of cells with and without ecDNA in a growing tumour population. We start with an analysis of neutral ecDNA dynamics. In this case the presence or absence of ecDNA does not affect cell fitness. In a tumour with *N* cells, the expected frequency of cells with ecDNA is

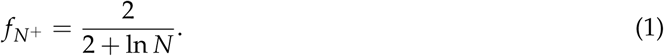

The frequency of cells with ecDNA 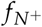 decreases with an increasing tumour population *N*. This might be surprising at first, given both cell populations have same fitness (Figure 2). However, while cells can lose ecDNA *N*^+^ → *N*^−^, the reverse is not possible 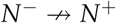. This asymmetry is sufficient to suppress cells with neutral ecDNA in large and growing cell populations. This has important consequences on the interpretation of *in vitro* ecDNA experiments, as the frequency of cells with ecDNA continuously declines under neutral dynamics. For fitness inferences it will be neccessary to show significant deviations from the dynamics as expected by equation (1), e.g. the frequency of negatively selected ecDNA should drop faster.

**Figure 2.**
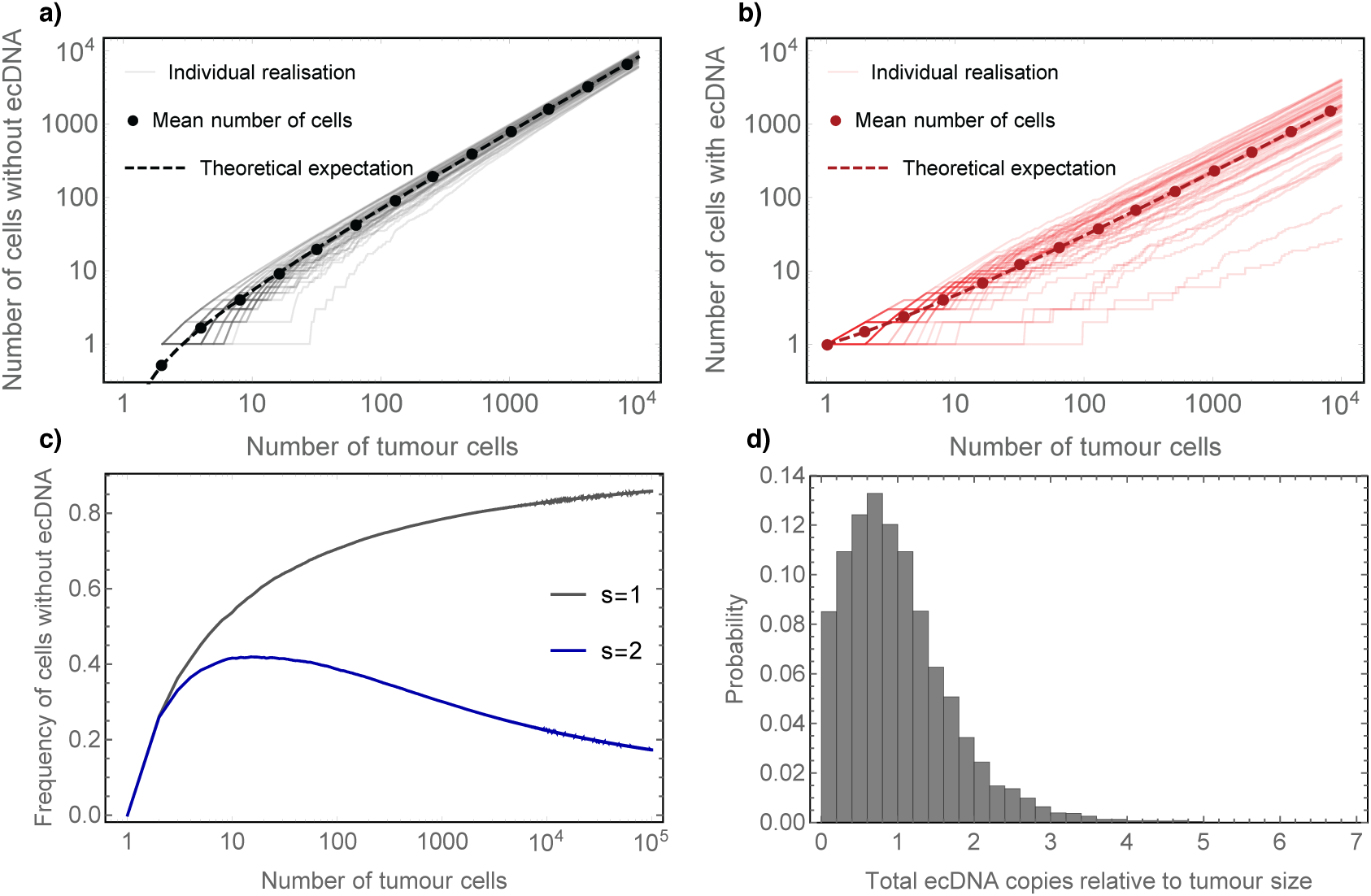
ecDNA population dynamics. Comparison of average deterministic dynamics of cells **a)** without and **b)** with copies of ecDNA. Dots show the average dynamics of neutral stochastic simulations, lines are individual realisation of the same neutral stochastic process and dashed lines show analytical predictions. Between tumour variation is considerable, especially for small tumour populations. **c)** Fraction of cells without ecDNA over time. In the neutral case *s* = 1 the tumour will be dominated by cells without ecDNA, also the fitness of cells with and without ecDNA is the same. Under strong positive selection, where cells with ecDNA have a selection advantage *s* = 2, the frequency of cells without ecDNA approaches 0. Even for strong positive selection we observe a transient increase of cells without ecDNA. **d)** Ratio of total number of ecDNA copies and total number of tumour cells for 10000 independent repeats of the same neutral evolutionary process. In many realisations there are as many or more ecDNA copies than tumour cells.

This does not imply that neutral ecDNA elements are not detectable in tumours at diagnosis. Based on equation (1) a tumour of 1 billion cells (10^9^) would be composed to 91% of cells without and 9% of cells with ecDNA. The ratio only changes marginally to 93% and 7% for a tumour with 10^12^ cells, which is within the detection threshold of current sequencing approaches. In Figure 2 we show examples of the ecDNA dynamics. The average dynamics of cells with or without ecDNA in a tumour population is captured exactly by equation (1), but stochastic differences between individuals are considerable. Importantly, although the number of cells carrying ecDNA is relatively small in a tumour of detectable size, the total number of ecDNA copies is of the same order of all tumour cells (Figure 2 c&d). This is of particular importance for genes involved in resistance evolution. Highly amplified copies of EGFR are a common resistance mechanism in targeted therapies [30, 31, 32]. Neutral ecDNA dynamics allows for single cells to carry many copies of ecDNA, enabling crowding effects of resistance mechanisms in small cell populations, e.g. single cells have a higher chance to acquire multiple resistance mechanisms. Consequently, the probability of pre-existing multi-drug resistance for genes amplified on ecDNA may be dramatically increased compared to chromosomal resistance mechanisms.

The deterministic dynamics changes once cells with ecDNA are under positive selection. Large tumours are ultimately dominated by cells with ecDNA (Figure 2c). However, even if ecDNA conveys a very strong positive selection advantage, the dynamics remains unintuitive. The frequency of positively selected cells with ecDNA changes approximately as

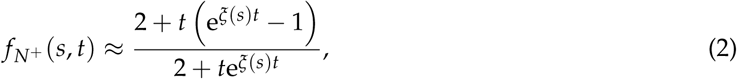

where the exponent *ξ*(*s*) is a function of the fitness coefficient *s*. Although large tumours are ultimately dominated by cells with ecDNA, cells without ecDNA initially rise in frequency and are only later outcompeted (Figure 2c). This transient effect is probably unobservable in large tumour populations, but may be important for cell line, organoid or xenograft experiments, urging a cautious interpretation of ecDNA fitness assays. For example, an initial decline of the frequency of cells with ecDNA can indicate neutral fitness, but also can be consistent with a fitness advantage. A robust experimental inference of ecDNA fitness will require growth to sufficiently large cell populations. Furthermore, even under very strong positive selection our model predicts the frequency of cells without ecDNA to always be above zero and thus one expects a mixed tumour population at any stage of the evolutionary process. As a consequence, the notion of clonal versus sub-clonal ecDNA copies becomes ambiguous [33, 34, 35], making it more difficult to determine if ecDNA was a founder event or occurred later during tumour evolution.

### Inter- and intra-tumour heterogeneity of ecDNA

We now turn to the stochastic properties of the ecDNA dynamics. Primarily, we want to quantify the expected inter- and intra-tumour heterogeneity. We therefore change from a deterministic two population model to a demographic description of ecDNA dynamics, which tracks the number of ecDNA copies in each cell at any given time (see Methods for details). This results in an infinite set of coupled differential equations, for which a fully closed analytical solution is challenging. However, we can derive important approximate properties of the time dependent probability distribution. For neutral ecDNA copies, we find that the moments of the ecDNA copy number distribution in tumour cells change in time according to

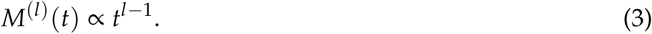

In particular, this implies that the mean number 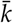 of neutral ecDNA copies remains constant 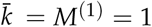, whereas the variance *σ*^2^ of the ecDNA distribution increases linearly in time *σ*^2^ = *M*^(2)^ − 1 = *t*. This also explains why under neutral dynamics the expected number of ecDNA copies in a growing tumour population is approximately the same as the number of tumour cells, despite the declining frequency of cells with ecDNA. In addition, this implies that on average neutral ecDNA amplifications would appear copy number neutral in a standard bioinformatics Copy Number Variation (CNV) pipeline for sufficiently large tumour bulks samples. Thus, neutral ecDNA amplifications may be more common than generally appreciated [3]. Novel bioinformatic tools to discover circular DNA structures help to solve some of these problems [3, 36]. Moreover, the polynomial scaling of all higher moments shows that inter-tumour ecDNA heterogeneity is expected to increase with time.

If cells with ecDNA have a fitness advantage, the moments also have a polynomial time dependence

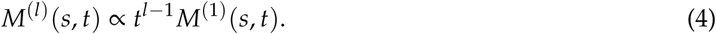

However, now the first moment is not constant but increases in time *t* depending on the selection coefficient *s*, see Methods. This difference compared to the expected constant mean number of neutral ecDNA copies is important for fitness inferences. Correspondingly, measuring the change of the first moments (e.g. mean and variance) of the ecDNA distribution with time can in principal inform about the (non)-neutral nature of ecDNA amplifictions. However, this would require high resolution temporally resolved single cell information of ecDNA copies, which is sparse in current clinical practice, but is available in *in vitro* experiments. With the fast development of single cell sequencing technology, a moment based method for ecDNA fitness inference may become more suitable for clinical settings in near future.

The lack of time resolved information on tumour evolution necessitates alternative evolutionary measures. One possibility is to quantify the expected distribution of ecDNA copies among (many) single cells of tumours, which is inferable from single time data and thus is more suitable to test ecDNA evolutionary dynamics in clinical practice. This is motivated by mutational site-frequency based methods in species and tumour evolution [10, 29]. There is an important distinction. The mutational site frequency spectrum quantifies cell fractions of point mutations, whereas ecDNA copy number distributions correspond to the number of cells with a certain number of ecDNA amplifications. We find that the number of cells *N*_*k*_ with *k* ecDNA copies is expected to scale as

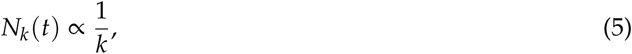

i.e. inversely with the number of ecDNA copies. Surprisingly, for a constant fitness coefficient *s* (e.g. no dosing effects of ecDNA), the dynamics among cells with ecDNA is again neutral and thus the scaling of the ecDNA copy distribution is again given by (5). Based on equation (5) a tumour should contain many cells with few ecDNA copies and fewer cells with many ecDNA copies (Figure 4). Stochastic simulations confirm the theoretical prediction even for small tumours (e.g. *N* = 10^4^ in Figure 4).

**Figure 3.**
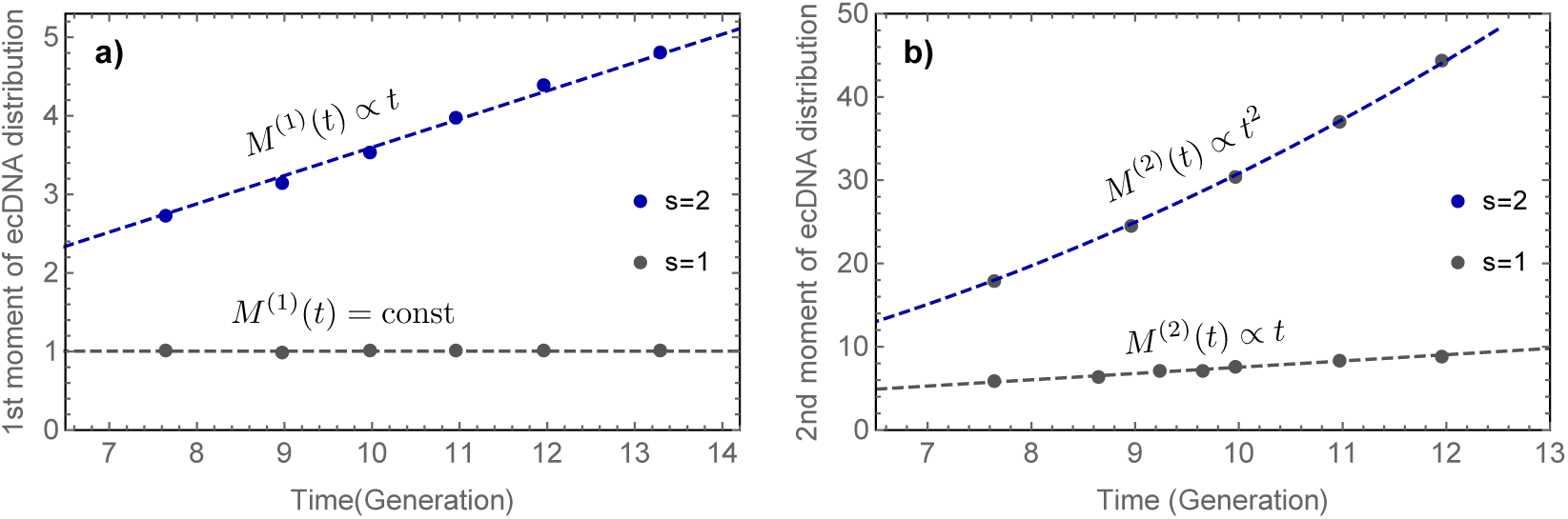
a) First and b) second moment of the ecDNA copy number distribution. In the neutral case (*s* = 1, grey) the mean number of ecDNA copies remains constant and the variance increases linearly in time. Under positive selection (*s* = 2, blue) the mean number of ecDNA copies increases in time. Stochastic simulations (points) are in very good agreement to theoretical predictions of polynomial increasing moments with time (dashed lines), see Eqs. (3) and (4).

**Figure 4.**
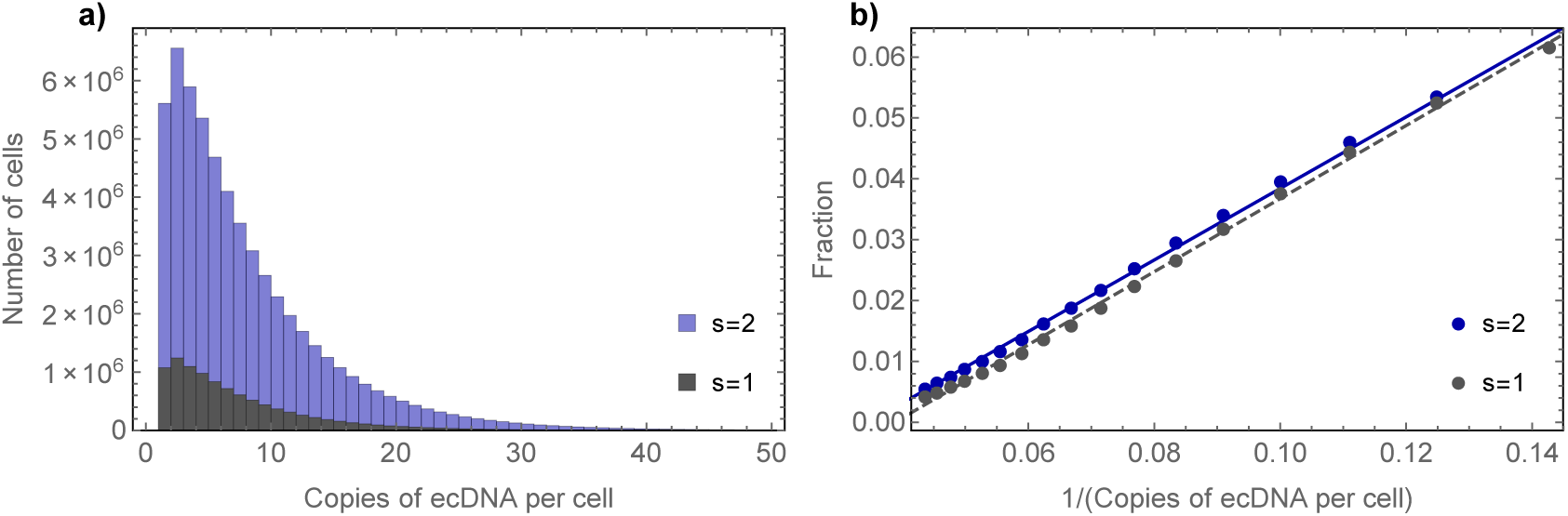
Distribution and scaling of the ecDNA copy number distribution. **a)** Distribution of the ecDNA copy number distribution for neutral (grey) and positively selected (blue) ecDNA evolution for 1000 repeats of stochastic simulations for tumours of 104 cells. Overall more cells carry copies of ecDNA if positively selected compared to the neutral case. **b)** The scaling of the ecDNA distribution follows the predicted 1/*k* scaling (dots = stochastic simulations, lines = theoretical expectation). However, the scaling law is the same for neutral evolution and constant positive selection and cannot discriminate neutrality from constant positive selection.

The difference to the neutral power law scaling of chromosomal point mutations is noteworthy. In an exponentially growing tumour population, point mutations scale as *f* ^−2^ with mutation frequency *f* [13, 14, 10], whereas the ecDNA copy distribution scales as *k*^−1^ with the number of ecDNA copies *k*. In addition, the scaling of the ecDNA copy number distribution alone cannot distinguish neutrality and selection. This is different for point mutations, where the site frequency spectrum can in principal distinguish neutrality from sufficiently strong selection [29, 37]. The scaling of the ecDNA copy number distribution depends solely on the within dynamics of cells with ecDNA. While the scaling of the ecDNA copy number distribution is insufficient to distinguish neutral and constant positive selection, it can distinguish those cases from a scenario where cells with different copy numbers of ecDNA have different fitnesses.

### Shannon diversity of ecDNA distributions

A third option for an ecDNA fitness inference is to combine properties of the intra-tumour ecDNA heterogeneity and time series information. One promising candidate is the Shannon diversity index *H* of a tumour population with regard to the ecDNA distribution. We find that the Shannon diversity depends on both, the strength of selection (fitness coefficient *s*) and the total time selection shapes the composition of the tumour population. The Shannon diversity index *H* changes approximately as

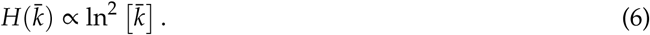

This predicts a quadratic logarithmic dependence of the Shannon diversity index *H* and the mean number of ecDNA elements 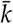 per cell. This dependence is recapitulated in stochastic individual based simulations. For numerical agreements, as shown in Figure 5a, we fit a parametrised curve 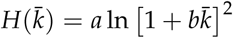 to stochastic simulations, using approximate bayesian computation (see Methods for details).

**Figure 5.**
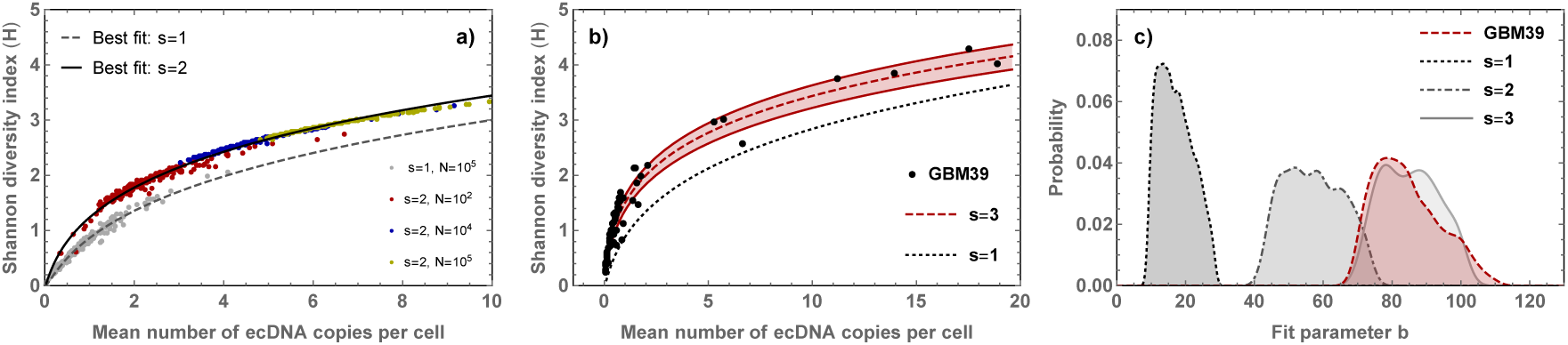
Circular extra-chromosomal *EGFRvIII* amplification in GBM39 is under strong positive selection. **a)** Shown is the Shannon diversity index *H* for stochastic simulations (each dot is a single realisation of the stochastic process) for varying degrees of selection and final sizes of the tumour population. For different fitness coefficients *s*, the Shannon diversity index falls on a fixed contour (e.g. dashed line: *s* = 1, black line: *s* = 2). Furthermore, the Shannon diversity increases towards higher values of *H* and 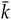 if ecDNA is positively selected but remains clustered around 1 in the neutral scenario. **b)** Shannon diversity index for 72 repeats of *in vitro* growth experiment using the glioblastoma cell line GBM39 (black dots), data from [1]. The Shannon diversity follows a quadratic logarithmic contour as predicted by theory. The dashed line shows the best fit for a fitness advantage of *s* = 3, shaded area corresponds to 95% credibility intervals. Neutral ecDNA dynamics (black dashed line) does not describe the observed patterns of Shannon diversity in the GBM39 cell line. **c)** Posterior distributions of approximate bayesian fits of the parametric curve 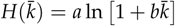 to stochastic simulations under varying selection strength *s* (gray lines) and GBM39 cell line experiments (red). The ABC suggests very strong positive selection for cells with ecDNA in the GBM39 cell line (*s* = 3). Neutral ecDNA dynamics (*s* = 1, dotted line) does not describe the observed diversity.

For neutral ecDNA copies, simulations cluster at small Shannon diversity and small mean number of copies of ecDNA per cell. Interestingly, if cells with ecDNA have a fitness advantage *s*, although the logarithmic dependence is maintained, there are two major differences from the neural case. Firstly, the curve shifts to higher diversity values compared to the neutral case. Secondly, the diversity within a tumour of positively selected ecDNA increases in time and travels along sharply defined contours for constant fitness coefficients *s* (Figure 5a). This is a direct consequence of the linearly increasing mean number of ecDNA copies (Figure 3a). These two properties of the Shannon diversity index suggest a principal strategy to distinguish neutral from positive selection from single time point measurements. For neutral evolution, individual realisations of the stochastic process fall onto a uniquely parametrised curve 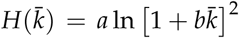 that clusters around a mean ecDNA copy number of 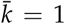. In contrast, positive selection of ecDNA leads to significant deviations form the neutral contour, shifting *H* to both higher Shannon diversity and higher average ecDNA copy numbers (Figure 5).

We test this theoretical prediction of the fitness effect of ecDNA on the Shannon diversity in repeated growth experiments of a glioblastoma cell line. In a recent publication Kirsten Turner and colleagues quantified the ecDNA copy number distribution in an *in vitro* growth experiment using the glioblastoma cell line GBM39 with circular extra-chromosomal *EGFRvIII* amplification [1]. In total, 72 repeats of the same cell line experiment were done using low pass (median coverage: 1.19x) whole genome sequencing data. This allowed us to quantify the contour of the Shannon diversity *H* for these experimental repeats (Figure 5b). We find a striking consistency of theoretical expectations and experimental observations. The cell line experiment shows a quadratic logarithmic dependence between the Shannon diversity and the average copy number of *EGFRvIII* (ecDNA element) for all experimental repeats. Even more striking, the Shannon diversity index of the circular extra-chromosomal *EGFRvIII* amplification changes along a single contour, suggesting a maintained fitness effect of the *EGFRvIII* amplification across experimental repeats. The curve is shifted towards higher Shannon diversity compared to the neutral expectation, suggesting that the extra chromosomal *EGFRvIII* amplification conveys a considerable fitness advantage in the GBM39 glioblastoma cell line. Figure 5c shows approximate bayesian fits (ABC) for different fitness scenarios in stochastic computer simulations and best inferences for the GBM39 cell line. The ABC inference suggests an almost 300% fitness increase (*s* = 3) for cells with compared to cells without the *EGFRvIII* amplification. The resulting parametric curve for *s* = 3 is shown in Figure 5b (dashed red line). It recovers the experimental Shannon diversity extremely well. In contrast, neutral ecDNA dynamics fails to describe the diversity patterns of ecDNA (black dotted line).

## Discussion

Here we developed and analysed theoretical models of ecDNA population dynamics in a growing tumour. The random segregation of ecDNA causes surprisingly unintuitive dynamics that can be markedly different compared to the dynamics of chromosomal point mutations. Consequently, common methods to quantify somatic evolutionary processes such as phylogenetic trees, dN/dS, or mutational site frequency spectra are not directly applicable to quantify ecDNA evolutionary dynamics. However, as we have shown here, evolutionary models of ecDNA dynamics can be constructed and analysed. Our analysis points to a very high fitness advantage (300%) of the *EGFRvIII* amplification in the GBM39 cell line. This is consistent with recent functional evidence of the *EGFRvIII* amplification [1, 9], as well as clinical evidence of poor survival of patients with ecDNA amplifications [38]. It will be highly interesting to quantify evolutionary histories of ecDNA in individual patients in future studies.

Our theoretical model of ecDNA population dynamics is of course build on assumptions. We assume that ecDNA segregation is random and ecDNA copies are amplified (doubled) prior cell divisions. Most importantly, we assume constant fitness. Dosed ecDNA fitness effects and their impact on ecDNA diversity requires a more detailed analysis in follow up studies. Yet, current experimental observations seem to support such a minimal evolutionary model. Larger experimental and/or clinical studies may unearth observations that will require us to consider more involved evolutionary scenarios.

## Acknowledgments

We are very grateful to Vineet Bafna and Paul S. Mischel for sharing data on the distribution of ecDNA copy numbers in the GBM39 cell line growth experiments. B.W. is supported by the Barts Cancer Charity.

## Methods

### Mathematical description of ecDNA dynamics

#### A deterministic two population model without selection

In the simplest representation of the model, we discriminate cells that do or do not carry copies of ecDNA. We denote cells with copies of ecDNA as *N*^+^(*t*) and cells without copies of ecDNA with *N*^−^(*t*). We can write for the change of these cells in time *t*

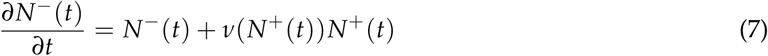

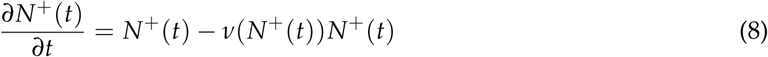

Cells with ecDNA *N*^+^ can lose all copies of ecDNA, but the reverse is not possible. ecDNA can not gained back once lost. The function *ν*(*N*^+^)(*t*) corresponds to the at the moment undetermined loss rate of ecDNA due to random complete asymmetric segregation, e.g. one daughter cell with *k* and the other daughter cell with 0 ecDNA copies. We now look at the fraction of cells that do not carry ecDNA 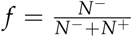. It follows that

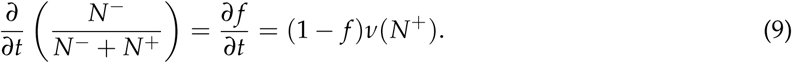

Again, for now both *f* and *ν* are unknown functions. However, from the equation above we can also write

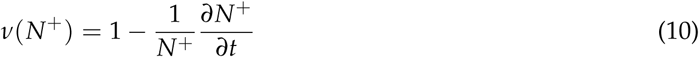

and thus it follows

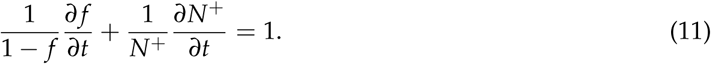

This equation can be integrated and after rearranging terms, we find the formal solution for *N*^+^(*t*),

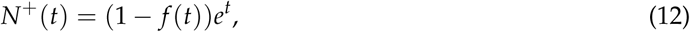

with initial conditions *N*^+^(0) = 1 and *f* (0) = 0. Assuming the tumour grows exponentially over time, e.g. *N*^−^(*t*) + *N*^+^(*t*) = *e*^*t*^, we get an expression for the number of cells without ecDNA

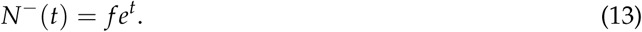

Stochastic simulations show that 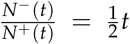. It follows that the fraction of cells without ecDNA elements in the neutral case changes as

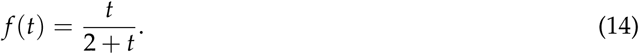

in time *t* (time measured in generations). This also allows us to find an exact expression for the expected loss of ecDNA copies *ν*(*N*^+^) within the subpopulation of cells with ecDNA *N*^+^. Returning to equation (10) we find

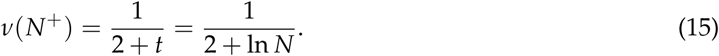

The rate of novel production of cells without ecDNA *ν* reduces with tumour size. However, the reduction is logarithmic in tumour size *N* and thus approaches 0 only slowly. Even in very large tumours, the rate is considerable, e.g. *ν*(*N* = 10^12^) = 0.034. Given 10^12^ cells this would translate to *N*^+^ → *N*^−^ ≈ 6 × 10^10^ novel cells without ecDNA emerging from cells carrying ecDNA by chance due to uneven ecDNA inheritance.

#### A deterministic two population model with selection

How will the dynamics change if cells with ecDNA are under positive selection? We therefore introduce a selection coefficient *s*, where *s* = 1 corresponds to the neutral case and *s* > 1 implies a fitness advantage. This leads to a modification of above equations and they take the form:

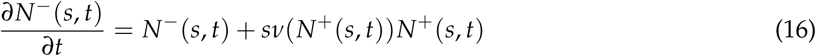

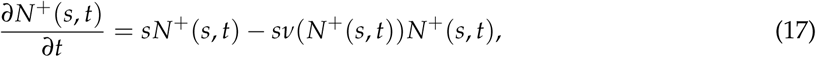

With the same procedure as above we can derive a general expression that is independent of *ν*(*N*^+^),which becomes

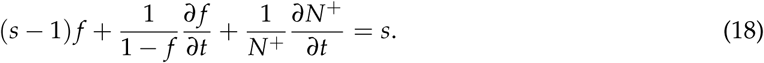

Note, in the neutral case *s* = 1 this reduces to equation (11). This expression can be solved and we can write

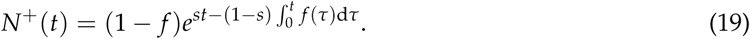

Stochastic simulations suggest a power law dependence of the form

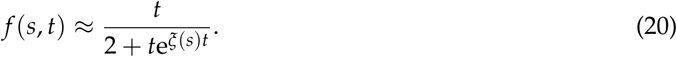

Hence, the fraction of cell with copies of ecDNA is

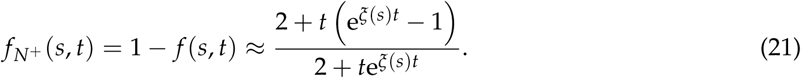

#### Stochastic dynamics of ecDNA elements

We now discuss the stochastic dynamics of ecDNA elements without selection. We therefore implement a demographic model of ecDNA, where we follow the number of cells *N*_*i*_(*t*) that carry *i* copies of ecDNA. Upon proliferation these copies are amplified and 2*i* copies are randomly distributed between both daughter cells. The number of copies *k* in a daughter cell then follows a Binomial distribution with success rate 1/2

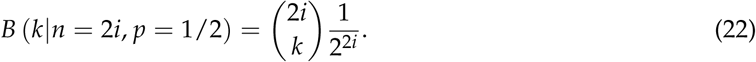

This allows us to write for the demographic model of the number of cells *N*_*k*_ with *k* copies of ecDNA

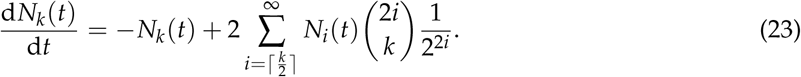

We can decouple population growth and demographic changes via the substitution *N*_*i*_(*t*) = *M*(*t*)*ρ*_*i*_(*t*) such that *ρ*_*i*_ is the density with 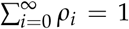 and *M*(*t*) corresponds to the population growth. In our case as a sanity check we can write

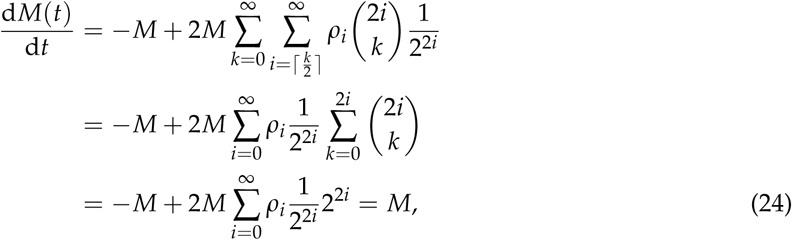

and the total cell population does growth exponentially *M*(*t*) = *M*(0)e^*t*^. This allows us to rewrite equation (23) for the densities *ρ*_*k*_, which gives

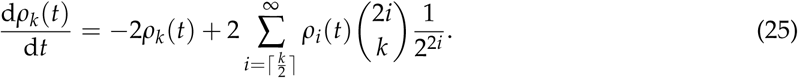

#### Moments dynamics

Equation (25) enables us to discuss the Moments dynamics of the ecDNA copy number distribution. In general, the *l*-th moment of the distribution is defined as

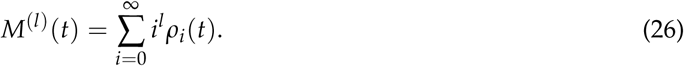

We find that the first three moments of the distribution can be calculated exactly. We furthermore give approximate scaling expressions for all higher moments of the distribution. *M*^(0)^ is just the sum over the density and by definition constant. The first moment corresponds to the average number of ecDNA copies per cell and we can write

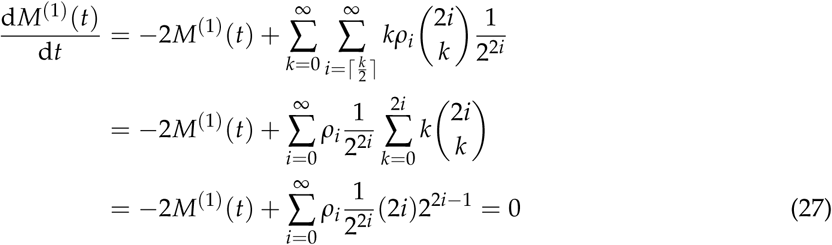

We therefore find *M*^(1)^(*t*) = const and the constant is given by *M*^(1)^(0), which in most cases discussed here is 1. For the second moment we find

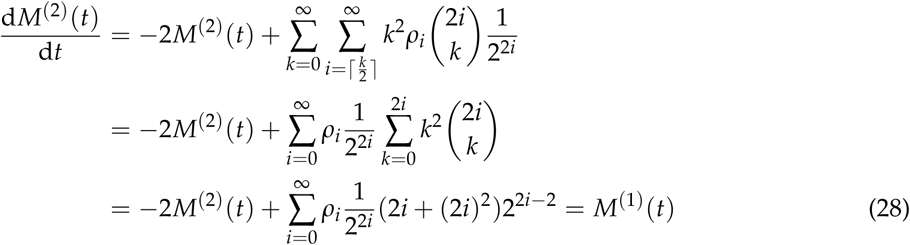

As we have *M*^(1)^ = 1, it follows that the second moment *M*^(2)^ = *t* increases linear in time. The general case for the *l*-th moment leads to a differential equation of the form

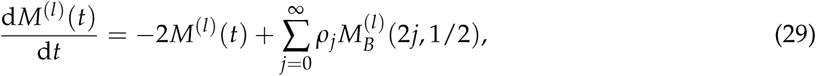

where 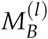 is the *l*-th moment of the Binomial distribution. There are no general closed expressions for the higher moments of the Binomial distribution. Approximate expressions can be obtained using the moment generating function

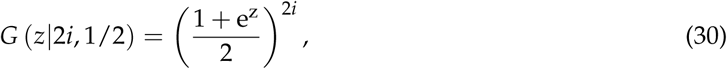

and the moments then become

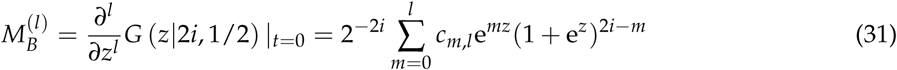

and therefore 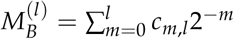. To get expressions for the coefficients *c*_*m*,*l*_ we can differentiate the moments equation one more time and get the recursion relation

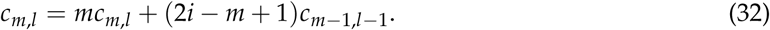

For example, 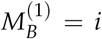 and 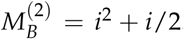. The moments of 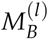 are polynomials over *i* and once plugged back into equation (29) they become a linear combination of moments *M*^(*i*)^ of the ecDNA copy number distribution *ρ*. The maximal possible rank of the *l*-th moment emerging in equation (29) is given by the maximal degree of the polynomial in 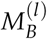 which is given by the coefficient *c*_*l*,*l*_ in above recursion. In this case, the recursion takes the form

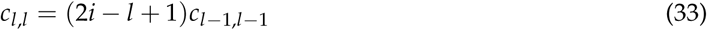

which has the solution 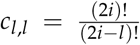. Keeping only the coefficient with the highest power, we get 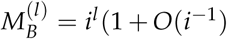. Returning to equation (29) and using the approximate expression for 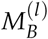 we get

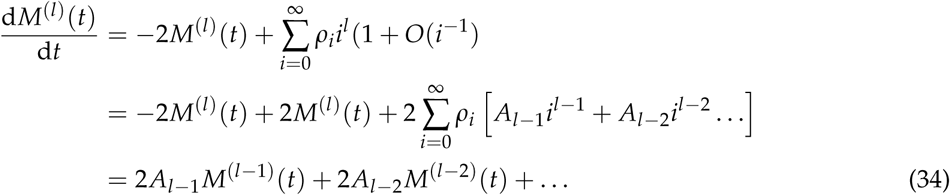

The derivative of the *l*-th moment depends linearly on all lower moments. Unfortunately, finding closed solutions for all coefficients *A* _*l*−*k*_ appears c hallenging. However, the general scaling behaviour becomes clear, and we have

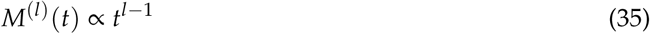

In the neutral case, the first moment remains constant and all higher moments diverge polynomial in time. This is in agreement with individual based stochastic simulations of the ecDNA dynamics Figure 3.

#### Continues limit and scaling wave solution

In the following we are interested in the scaling behaviour of the ecDNA copy number distribution. The general equation (25) considers discrete copy number states. To make further analytical progress, we consider continues states in the following calculations. Such an approximation likely describes the case of many ecDNA copies well, but might be inaccurate for cells with very few ecDNA copies. Under the continues assumption, the change of the ecDNA distribution becomes

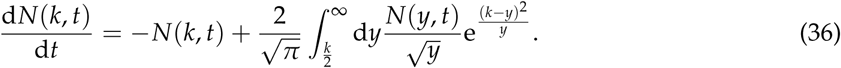

Where *N*(*k, t*) is the number of cells with ecDNA copy number *k* at time *t* and the Binomial distribution in equation (25) was replaced by a Normal distribution assuming a limit of large *k*. Similarly, we can write for the density

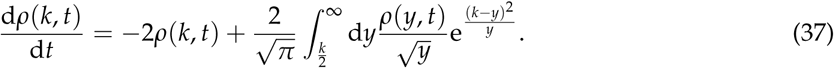

Given the exponential character of the ecDNA distribution, we proceed with an ansatz in the form of a scaling wave solution

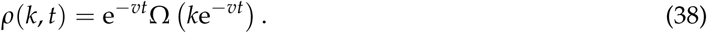

Plugging Ansatz (38) back into equation (37) and setting *k*/2 → 0, we get

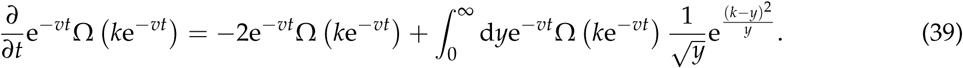

The left side of equation (39) then becomes

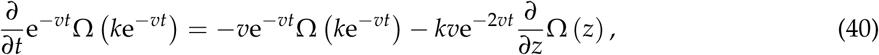

with *z* = *k*e^−*vt*^. For *v* = 2 the first terms on the right hand side of equation (39) and (40) cancel and equation (39) becomes

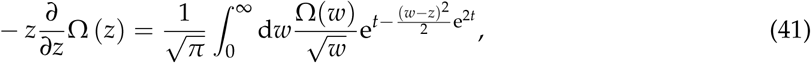

where as above *z* = *k*e^−2*t*^ and *w* = *y*e^−2*t*^. With a final substitution *m* = (*w* − *z*)e^*t*^, substituting *m* for *w* and taking the limit *t* → ∞ we get

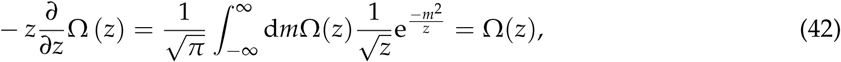

with the solution

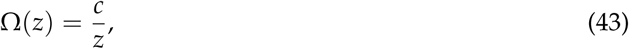

and an undetermined integration constant *c*. Finally this gives for the scaling behaviour of the ecDNA copy number distribution

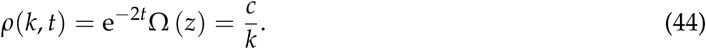

This predicts that the number of cells with *k* ecDNA copies scales inversely with *k*. We find the same scaling behaviour in individual based stochastic simulations of the ecDNA copy number distribution, Figure 4.

#### Dynamics of ecDNA under constant positive selection

Here, we consider the situation, where cells containing ecDNA grow with rate *s* > 1. Then, Eq. (23) becomes

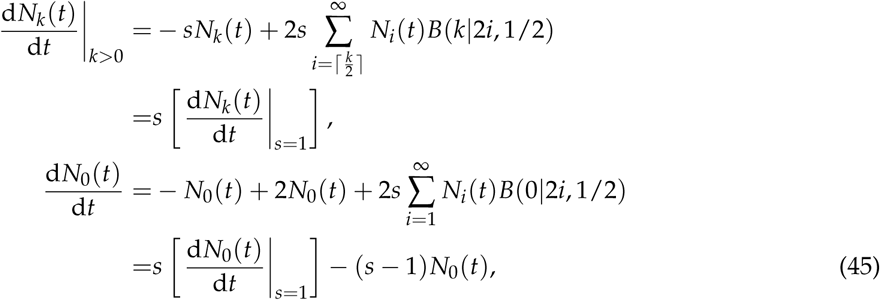

Again, we decouple population growth and demographic changes via the substitution *N*_*i*_(*t*) = *M*(*t*)*ρ*_*i*_(*t*) such that *ρ*_*i*_ is the density with 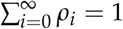 and *M*(*t*) corresponds to the population growth. Similar to Eqs. (24) and (25), we get

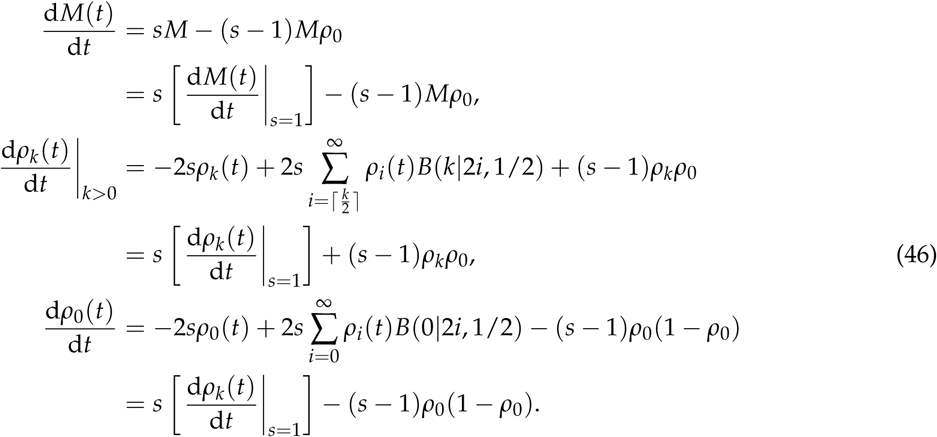

In the case of constant positive selection, the demographic dynamics gains additional non-linear terms compared to the neutral case.

Taking into account these terms, we can find new moments dynamics:

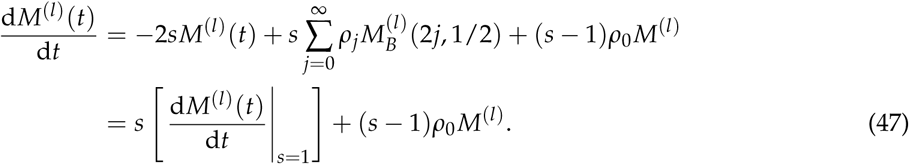

Hence,

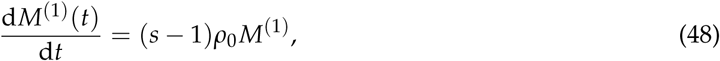

and as a result,

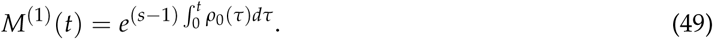

Similarly,

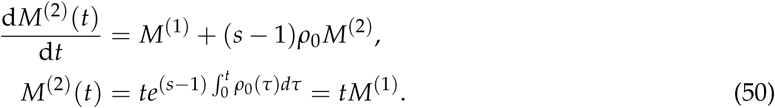

Similar to Eq. (34), we can show that

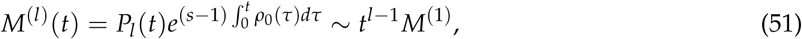

where *P*_*l*_ (*t*) is some polynomial over *t* of degree *l* − 1. Therefore, the power law of the moment dynamics with incrementing powers observed for neutral selection is also preserved in the constant selection case. However, closed expressions are more challenging, as the equation for *M*^(1)^ includes additional terms.

#### Shannon diversity index

For the Shannon diversity index *H* we can write

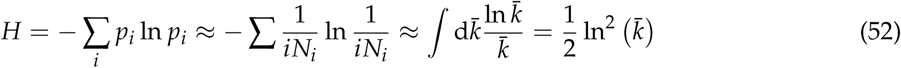

The first approximation considers the scaling of the ecDNA copy distribution followed by approximating the sum by integration. We therefore find

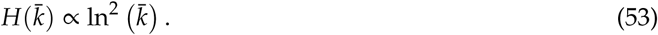

Stochastic computer simulations recapitulate the quadratic logarithmic scaling of the Shannon diversity index (Figure 5a).

### Stochastic individual based simulations of ecDNA population dynamics

We implemented individual based stochastic computer simulations of the ecDNA population dynamics in C++. For each cell, the exact number of ecDNA copies is recorded trough the simulation. Cells are chosen randomly but proportional to fitness for proliferation. The proliferation rate of cells without ecDNA is set to *r*^−^ = 1 (time is measured in generations). A fitness effect for cells with ecDNA then corresponds to a proliferation rate *r*^+^ = *s*. Here *s* > 1 models a fitness advantage, 0 < *s* < 1 a fitness disadvantage and *s* = 1 corresponds to no fitness difference (neutral dynamics, *r*^+^ = *r*^−^). If a cell is chosen for proliferation, the number of ecDNA copies in that cell are doubled and randomly distributed into both daughter cells according to a Binomial trail with success rate 1/2. If a cell carries no ecDNA, no daughter cell inherits ecDNA. We terminate simulations at a specified cell population size. We output the copy number of ecDNA for each cell at the end of each simulation, which allows us to construct other quantities of interest, such as the ecDNA copy distribution, the time dynamics of the moments, or the Shannon diversity index *H*.

### Approximate Bayesian Computation

We implement Approximate Bayesian computation (ABC) to fit the Shannon diversity index *H* to stochastic simulations of ecDNA dynamics with different fitness strength *s*. We run 10000 simulations for a fixed fitness coefficient *s* to generate a distribution of Shannon diversity indices *H* in dependence of the mean copy number distribution 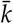 for each simulation. We then fit the parametric curve 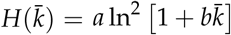 to each evolutionary scenario independently. Here *a* and *b* are free parameters fitted by the ABC algorithm. We therefore choose *a* and *b* randomly from uninformed uniform distributions (*a* ∈ [0, 0.3] and *b* ∈ [0, 200]) and reject values for which the coefficient of determination 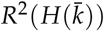 of the parametric curve with random parameters *a* and *b* and the simulated data set is *R*^2^ < 0.995. This gives a posterior distribution for the parameters *a* and *b*. We do the same ABC procedure for the Shannon diversity distribution of the GBM39 cell line experiment, with a threshold of *R*^2^(*H*) < 0.93. We use the posterior parameter distributions to generate credibility intervals via bootstrapping.

